# Verapamil/Curcumin treatment attenuates the behavioral alterations observed in Williams Syndrome mice by regulation of MAPK pathway and Microglia overexpression

**DOI:** 10.1101/2021.02.01.429086

**Authors:** Paula Ortiz-Romero, Gustavo Egea, Luis A Pérez-Jurado, Victoria Campuzano

**Author notes:** **Correspondence:** Victoria Campuzano, ^1^Departament de Biomedicina, Facultat de Medicina i Ciències de la Salut, Universitat de Barcelona, 08036 Barcelona, Spain.

## Abstract

Williams-Beuren syndrome (WBS) is a rare neurodevelopmental disorder characterized by a distinctive cognitive phenotype for which there currently are not any effective treatments. We investigated the progression of behavioral deficits present in CD (complete deletion) mice, a rodent model of WBS, after chronic treatment with curcumin, verapamil and a combination of both. These compounds have been proven to have beneficial effects over different cognitive aspects of various murine models and thus, may have neuroprotective effects in WBS. Treatment was administered orally dissolved in drinking water. A set of behavioral tests demonstrated the efficiency of combinatorial treatment. Some histological and molecular analyses were performed to analyze the effects of treatment and its underlying mechanism in CD mice. Behavioral improvement correlates with the molecular recovery of several affected pathways regarding MAPK signaling, in tight relation with the control of synaptic transmission. Moreover, CD mice showed an increased activated microglia density in different brain regions, which was prevented by treatment. Therefore, results show that treatment prevented behavioral deficits by recovering altered gene expression in cortex of CD mice, reducing activated microglia and normalizing *Bdnf* expression levels. These findings unravel the mechanisms underlying the beneficial effects of this novel treatment on behavioral deficits observed in CD mice, and suggest that the combination of curcumin and verapamil could be a potential candidate to treat the cognitive impairments in WBS patients.

## 1 Introduction

Williams-Beuren syndrome (WBS, OMIM 194050) is a rare neurodevelopmental disorder with an estimated prevalence of 1 in 7500-20000 newborns that is caused by the heterozygous deletion of 26-28 contiguous genes (1.55-1.83 Mb) on chromosome 7q11.23 (Strømme et al., 2002; Bayés et al., 2003)Together with some cardiovascular features, WBS individuals present mild to moderate intellectual disability, with a mean intelligence quotient (IQ) that ranges between 50 and 60 (Martens et al., 2008). They present a unique cognitive phenotype that includes severe deficits in visuospatial construction and an increased sociability, together with preserved linguistic abilities (Bellugi et al., 2000; Doyle et al., 2004). Given the complexity of the cognitive profile of WBS patients they often require assistance of a mental health professional. However, pharmacological intervention is usually inaccurate and only targeted to treat anxiety (Green et al., 2012; Martens et al., 2012). For this reason, it is interesting to study any potential treatment that might improve the cognitive impairments observed in WBS.

The complete deletion (CD) mouse model presents a cognitive phenotype very similar to that seen in humans with WBS (Segura-Puimedon et al., 2014) and has been used to define possible therapeutic interventions (Borralleras et al., 2015; Ortiz-Romero et al., 2018). The cognitive characterization of this animal model has shown impairments in motor coordination, anxiety-like behavior and hypersociability (Segura-Puimedon et al., 2014). Moreover, histological analyses of the brain have revealed disorganization in certain brain areas as well as an abnormal morphology of dendritic spines (Borralleras et al., 2015; Ortiz-Romero et al., 2018).

In the last years, the study of the therapeutic effects of natural phenols has gained attention. Accumulated evidence has described that curcumin, the major constituent of turmeric *(Curcuma longa),* exerts a variety of pharmacological effects due to its antioxidant, anti-inflammatory and neuroprotective properties (Hay et al., 2019). Recent studies have reported positive effects of curcumin over different cognitive aspects such as anxiety-like behaviors, memory deficits and motor impairments of different murine models (Spinelli et al., 2015; Aubry et al., 2019; Sanei and Saberi-Demneh, 2019). Many studies have described that its effects in the behavioral phenotype of mice models are mediated by an upregulation of BDNF expression (Zhang et al., 2012; Franco-Robles et al., 2014; Nam et al., 2014). BDNF has been described as a crucial molecule for neural development and plasticity processes (von Bohlen und Halbach and von Bohlen und Halbach, 2018), and its mechanism of action is highly dependent on a proper maintenance of intracellular ionic homeostasis (Li et al., 1999; Numakawa et al., 2001). Moreover, it has also been described to prevent neuroinflammation by modulating pathways related to *Nrf2* and MAPK signaling (Jin et al., 2018).

Verapamil is a widely used medication and its mechanism of action involves mainly the blocking of voltage-dependent calcium channels (Elliott and Ram, 2011), but it has also been proven to directly bind and block voltage-gated potassium channels (Ko et al., 2010) and to inhibit drug efflux pump proteins like P-glycoprotein (Bellamy, 1996). Although it has been mainly studied for cardiovascular applications, it has also been associated with positive effects on anxiety and memory processing in murine models (Quartermain and Garcia DeSoria, 2001; Biala and Kruk, 2008).

Given the properties of both compounds, we decided to explore the effects of each compound and a combinatorial treatment on the cognitive phenotype of CD mice. Results showed that only combinatory treatment with curcumin and verapamil in CD mouse model of WBS improved motor coordination, which in turn correlated with the normalization of the MAPK signaling pathway with the concomitant reduction of activated microglia in different brain regions.

## 2 Materials and methods

### 2.1 Ethics statement

The local Committee of Ethical Animal Experimentation (CEEA-PRBB Protocol Number: VCU-17-0021; and University of Barcelona Protocol Number: Campuzano-153.19) and Generalitat de Catalunya (Protocol Number DAAM-9494) approved all animal procedures in accordance with the guidelines of the European Communities Directive 86/609/EEC. The PRBB has Animal Welfare Assurance (#A5388-01, Institutional Animal Care and Use Committee approval date 05/08/2009), granted by the Office of Laboratory Animal Welfare (OLAW) of the US National Institutes of Health.

### 2.2 Animals’ maintenance

CD mice, a WBS murine model that carries a 1.3 Mb heterozygous deletion harboring from *Gtf2i* to *Fkbp6,* were obtained as previously described (Segura-Puimedon et al., 2014). CD mice were crossed with Thy1-YFP transgenic mice (B6.Cg-Tg(Thy1-YFPH)2Jrs/J, Jackson Laboratory) (Feng et al., 2000) to label pyramidal neurons. All mice were maintained on 97% C57BL/6J background. Genomic DNA was extracted from mouse ear punch to perform the genotyping using MLPA/PCR and appropriate primers, as previously described (Borralleras et al., 2015). Animals were housed under standard conditions in a 12 h dark/light cycle with access to food and water/treatment *ad libitum.* We used four groups of treatment (vehicle, verapamil (VER), curcumin (CUR) and combined (VER+CUR)) per genotype (WT and CD) in male mice, with n=7-11 in all groups.

### 2.3 Treatment administration and intake

Considering that the higher dose of VER used in human patients is 480 mg/day, we established an approximately equivalent dose of 10 mg/kg/day for our treatment (considering average mouse weight 25 g). VER was dissolved in drinking water and kept at −21 °C until its use.

CUR was administered at a dose of 60 mg/kg/day as previously reported (Babu et al., 2015). It was freshly prepared once per week, from a curcumin extract (Super Bio-Curcumin®, from Life Extension®, USA, total curcumoids complex with essential oils of turmeric rhizome by HPLC 400 mg) dissolved in drinking water with 1% DMSO.

Vehicle control group consisted of WT and CD mice drinking water with 1% DMSO. Experimental groups consisted of: 1) WT and CD mice treated with VER only; 2) WT and CD mice treated with CUR only and 3) WT and CD mice treated with both compounds (VERCUR).

Respective treatments were started at 8 weeks-old (young mice). All animals received their treatments for 1 month before starting the behavioral tests and it was maintained for the whole duration of these procedures until sacrifice at 12-16 weeks-old (young adult).

The amount of drink per cage was quantified and normalized to the number of animals per cage (2 to 4) and to the time between each change (48 to 60 hours). Consumption was not significantly different between groups (F_2.005,30.08_ = 0.4796, *p* = 0.6242) (Supplemental Figure 1(A)). The consumption per day was reduced in CD animals (two-way ANOVA, effect of genotype F_1,41_ = 11.11, *p* = 0.0018). However, individual group comparisons revealed that only VER+CUR groups showed significant differences mainly due to an increase in the consumption in WT animals. WT and CD animals presented no differences for the rest of treatments (Supplemental Figure 1(B)).

In agreement with previous reports (Segura-Puimedon et al., 2014), CD mice had significantly lower body weights compared to WT mice (effect of genotype F_1,84_ = 146.4, *p*<0.0001), with a significant effect of treatment (F_3,84_ = 3.407, *p* = 0.0213) due to a difference between WT and VER-consuming WT animals. No differences were observed for the other groups of treatment. (Supplemental Figure 1(C)).

### 2.4 Behavioral tests

All the experiments were performed during the light phase of the dark/light cycle by researchers blind to the different experimental groups. There was a constant illumination of about <10 lux. All the apparatus were cleaned after each mouse.

#### 2.4.1 Social interaction test

The social interaction test was conducted in an open field as previously described (Borralleras et al., 2015). Briefly, an empty wire cup-container was placed in the center of the arena and each individual mouse was allowed to freely explore the arena for 5 minutes. During this time, the amount of time sniffing the empty container was measured. After this, an intruder mouse was placed in the container and, again, the amount of time nose-to-nose sniffing the intruder mouse was measured during 5 minutes.

#### 2.4.2 Marble-burying test

The marble-burying test was conducted as previously described (Ortiz-Romero et al., 2018). Briefly, a polycarbonate rat cage was filled with 5 cm depth of bedding and lightly tamped down. A regular pattern of 20 glass marbles (five rows of four marbles) was placed on the surface of the bedding prior to each test. Animals were tested individually, and the number of buried marbles (considered buried when >2/3 of the marble was covered) was counted every five minutes for a total time of 20 minutes.

#### 2.4.3 Rotarod test

The rotarod test was performed as previously described (Borralleras et al., 2015). Briefly, the test was conducted using a Rotarod LE8500 apparatus (Panlab, Harvard Apparatus). First, a period of training was performed in which animals were trained until they could stay on the rod for 120 seconds at the minimum speed (4 rpm). After 5 minutes of rest the test was started, which consisted in measuring the latency to fall off the rod in consecutive trials of 4, 10, 14, 19, 24, 34 and 40 rpm, with a maximum of 60 seconds per trial and a rest of 5 minutes between trials.

### 2.5 Histological preparation

Animals were perfused with 1X phosphate buffered saline (PBS) followed by 4% paraformaldehyde (PFA). Brains were removed and postfixed in 4% PFA for 24 h at 4 °C, in PBS for 24 h at 4 °C and, afterwards, crioprotected in 30% sucrose for 24 h at 4 °C.

#### 2.5.1 Immunofluorescence

30 μm-thick coronal brain sections were permeabilized with 0.3% Triton-X and incubated with NH4Cl 50 mM for 30 min at room temperature. Blocking was performed with a solution of 2% bovine serum albumin and 3% fetal bovine serum for 2 h at room temperature. Afterwards, sections were incubated overnight at 4 °C with primary antibodies [rabbit polyclonal anti-IBA1 (1:1000, Wako) and goat polyclonal anti-IL1β (1:500, R&DSystems)]. After washing, secondary antibodies [anti-rabbit 1:1500, anti-goat 1:500 (Invitrogen)] were incubated for 1 h at room temperature. Sections were mounted with Mowiol mounting medium.

For IBA1/IL1β quantification, 1024×1024 pixel confocal fluorescent image stacks from these brain sections were obtained with a TCS SP5 LEICA confocal microscope, using a X20 objective. We obtained pictures of CA1 hippocampus, motor cortex and somatosensory cortex regions. Image J software was used for the image quantification. N = 3 - 5 animals per group were analysed, and three brain slices were analysed per animal. Percentage of activated proinflammatory microglia was calculated by relativizing the number of IL1β positive cells to the number of IBA1 positive cells.

#### 2.5.2 Morphological evaluation of different brain areas

150 μm-thick serial coronal brain sections were collected on a glass slide and directly mounted with Mowiol.

For the quantification of the number of YFP+ neurons, 1360×1024 pixel images of CA1 hippocampus and motor cortex were obtained with an Olympus DP71 camera attached to an Olympus BX51 microscope with an Olympus U-RFL-T source of fluorescence at 10x magnification. Image J software was used for quantification. N = 3-5 animals per group were analyzed.

For spine density and length analyses, 1024×1024 pixel confocal fluorescent image stacks from these tissue sections were obtained with a TCS SP2 LEICA confocal microscope, using a X63 (zoom x5) oil immersion objective. We obtained pictures of dendritic segments of 15 - 30 μm from randomly selected neurons in CA1 hippocampus and motor cortex sections. Image J software was used for quantification. Spine counts included all type of dendritic protrusions. N = 3 - 4 animals per group were analyzed, and ten dendritic segments were analyzed per animal. Spine density was calculated by relativizing the total number of spines to the length of the analyzed dendrite. Spine length was measured from the base of the spine to the end of the head of the spine.

### 2.6 RNA extraction and gene expression analyses

Brains were removed and the frontal cortex was isolated, immediately frozen, and kept at −80 °C until use. RNA was extracted from the cerebral cortex using RNeasy Mini Kit (Qiagen) following manufacturer’s instructions, and mRNA quality was evaluated with BioAnalyzer (Agilent).

RNA from WT and CD mice (n=3 each) per experimental group (vehicle-treated and VER+CUR-treated) was used to prepare twelve mRNA libraries following the standard Illumina protocol. Libraries were sequenced at the Centre Nacional d’Anàlisi Genòmica (CNAG-CRG, Barcelona, Spain) facilities, using an Illumina HiSeq2000 platform to produce over 60 million paired-end reads (100 nucleotide length) per sample. Reads were mapped to the Ensembl GRCm38 Mus musculus genome provided by Illumina iGenomes. Mapping was performed with STAR/2.5.3a. After a correction by percentage of coefficient of variation (%CV<20), we discriminated 13941 mapped protein coding genes to be analysed.

Expression values were relativized according to the average expression of the WT animals for each gene. A gene was considered to be differentially expressed between two groups when the fold-change was <0.75 or >1.25, and with a *p*<0.05. Results were validated by the expression analysis of the genes included in the WBS critical region (WSCR), 20 of which are represented in the analysis (Supplemental Figure 2) and are known to present reduced expression in WBS mice models (Li et al., 2009; Segura-Puimedon et al., 2014; Kopp et al., 2019).

We used the CPDB (available at http://www.consensuspathdb.org/) to investigate the functional associations of genes found to be either down- or up-regulated.

### 2.7 Statistical analysis

All data are presented as mean ± SEM. one-way ANOVA and two- or three-way ANOVA with Dunnett or Sidak’s post hoc analyses test were used when needed. Values were considered significant when *p*<0.05. GraphPad Prism 8 software was used for obtaining all statistical tests and graphs. Heatmap was generated using heatmap.2 function of the *gplots* package in R.

## 3 Results

### 3.1 Only the combinatory treatment with curcumin and verapamil improves the motor coordination in CD mouse model of WBS

Motor coordination was evaluated 30 days after treatment in the rotarod test in WT and CD mice receiving VER, CUR, VERCUR or vehicle. Three-way ANOVA (rpm, genotype, treatment) revealed significant effects of genotype (F_1,261_ = 17.29; p<0.0001) and interaction between genotype and treatment (VERCUR) (F_1,261_ = 11.87; *p* = 0.0007) (Figure 1(A). Post hoc analyses showed significant motor coordination deficits in the vehicle-treated CD animals up to 19 rpm in comparison with WT animals (Supplemental Table 1) replicating previous results of basal groups (Segura-Puimedon et al., 2014). This effect was prevented by VERCUR treatment (Figure 1(A), Supplemental Table 1). Results showed neither single VER nor single CUR treatments had any significant effect on the performance of treated animals in this test (F_1,254_ = 0.5085; *p* = 0.4764 for VER; F_1,264_ = 1.947; *p* = 0.1641 for CUR)(Figure 1(B) and (C)). None of the treatments had any effect in the results of WT mice in this test (effect of treatment F_3,256_ = 1.918, *p* = 0.1272) (Supplemental Figure 3).

**Figure 1.**
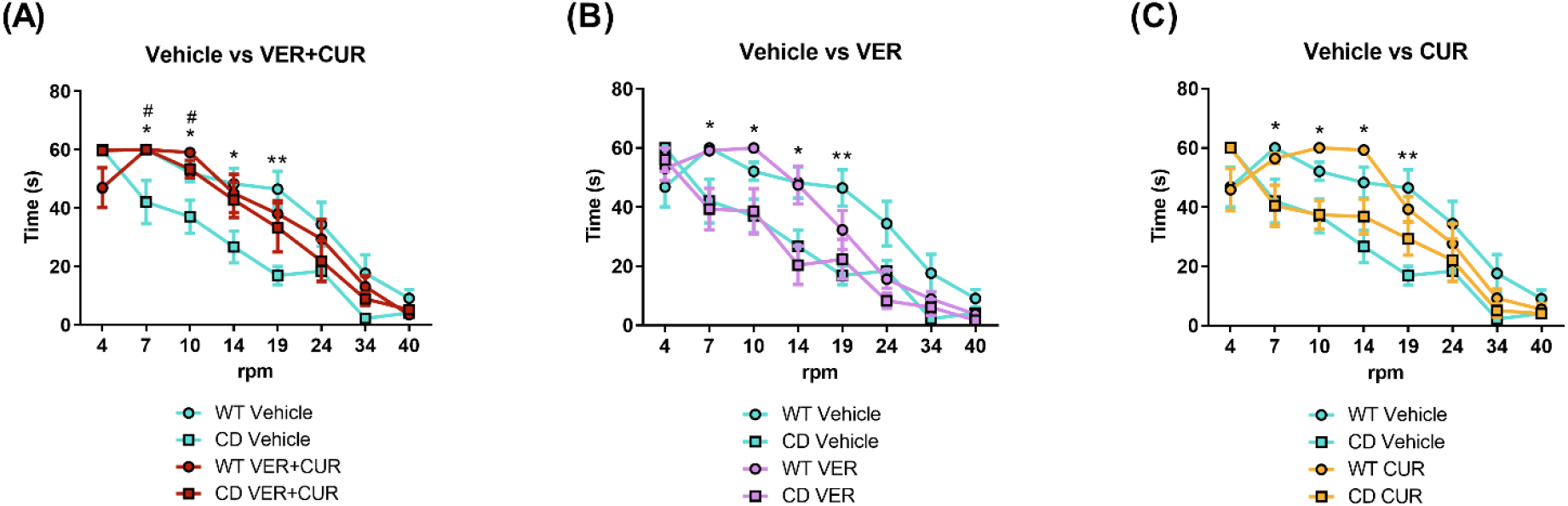
Motor coordination is improved in CD mice after treatment with verapamil and curcumin. **(A)** After combinatorial VER+CUR treatment CD animals performed better in the rotarod test (effect of treatment: F_1,261_=4.193; *p*=0.0416), staying in the rod for significantly more time than vehicle-treated CD mice (interaction between genotype and treatment: F_1,261_=11.87; *p*=0.0007). **(B)** Single treatment with VER did not have any effect on the performance of treated CD animals in the rotarod test (effect of treatment F_1,254_=3.031; *p*=0.0829; interaction between genotype and treatment F_1,254_=0.5085; *p*=0.4764). (C) Single treatment with CUR did not have any effect on the performance of treated CD animals in the rotarod test (effect of treatment F_1,262_=0.3435; *p*=0.5583; interaction between genotype and treatment F_1,262_=1.947; *p*=0.1641). Data are presented as mean±SEM of n=7-11 mice. Adjusted *p* values are shown with asterisks (genotype effect) or hashes (treatment effect) indicating values that are significantly different in individual group comparisons (two-way ANOVA, one for each rpm, followed by Tukey’s*post hoc* test). *,#*p*<0.05; **,##*p*<0.01

### 3.2 Combinatory treatment prevents hypersociability of CD mice

Social behavior was determined using the social interaction test. Three-way ANOVA (intruder, genotype, treatment) revealed significant effects of intruder in all the treatments (VER: F_1,62_ = 21.42; *p* < 0.0001; CUR: F_1,64_ = 4.017; *p* = 0.04; VERCUR: F_1,65_ = 14.41; *p* = 0.0003). In all cases, Sidak’s post hoc analyses revealed that the vehicle-treated CD group showed significant differences between the amount of time exploring the intruder mouse respect to the time exploring the empty cage (VER: *p* = 0.0025; CUR, *p* = 0.0087; VERCUR, *p* = 0.0013), replicating previous results (Segura-Puimedon et al., 2014) (Figure 2). This effect was prevented by VERCUR treatment *(p* = 0.5728) (Figure 2A), but not by VER treatment in which treated animals replicated the behavior of vehicle-treated mice, showing significant differences between the amount of time exploring the intruder mouse respect to the time exploring the empty cage (*p* = 0.0058) (Figure 2B),. In CUR treatment, the three way ANOVA analysis revealed a significant effect of treatment (CUR: F_1,64_ = 9.232; *p* = 0.0034) due to an increased interest for the empty cage regardless of genotype (Figure 2C)

**Figure 2.**
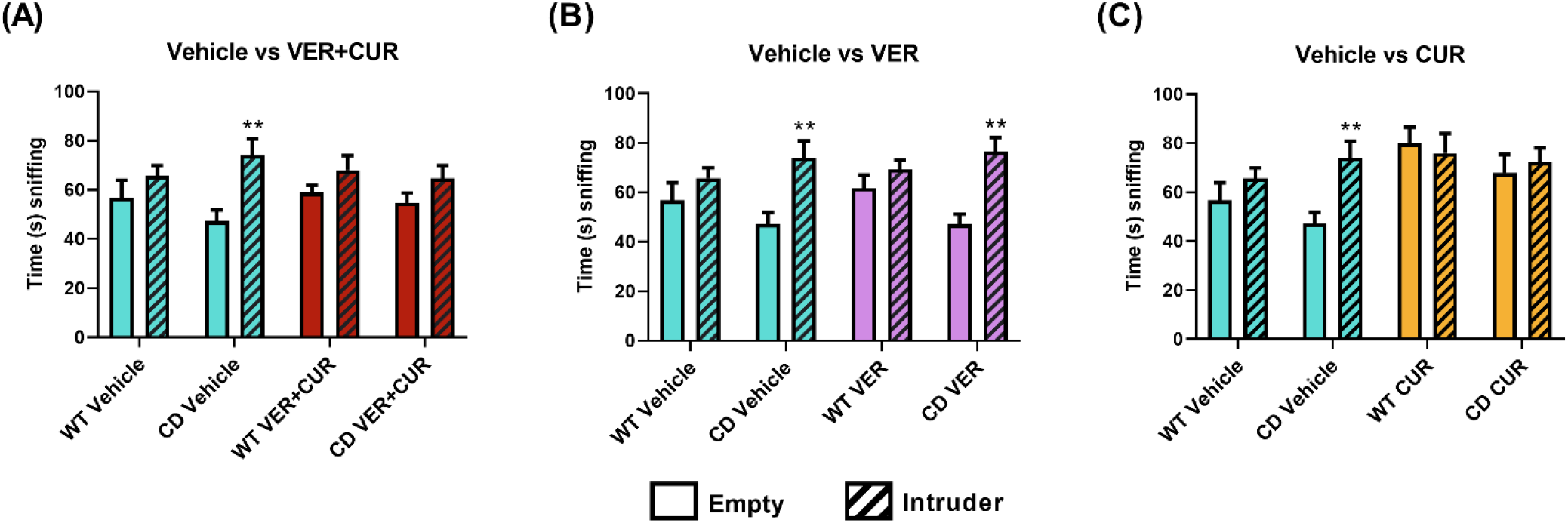
Hypersociability of CD mice is prevented after treatment with verapamil and curcumin. **(A)** In a direct social test, vehicle-treated CD animals presented an increased interest towards an intruder mouse in comparison with the empty container (*p*=0.0025), while VER+CUR-treated CD animals showed less interest in the intruder animal, exploring it the same amount of time as the empty container (*p*=0.5728). **(B)** Single VER treatment did not have any effect on the performance of CD animals in this test, as they presented an increased exploratory interest towards an intruder mouse as seen in vehicle-treated animals (*p*=0.0058). **(C)** Single CUR-treated CD animals did not show a significant interest towards the intruder animals (*p*=0.9830), but they also presented a tendency of increased interest towards the empty cage, which was also observed in WT animals. Data are presented as mean±SEM of n=6-10 mice. *p* values are shown with asterisks indicating values that are significantly different in three-way ANOVA with Sidak’s*post hoc* test. \**p*<0.05, \*\**p*<0.01.

### 3.3 Anxiety-like behavior of CD animals is not improved by any treatment

The evaluation of anxiety-like behavior was performed by counting the number of marbles buried in the marble burying test (MBT) after 1 month of treatment. Two-way ANOVA (genotype, treatment) revealed significant effects of genotype (F_1,66_ = 160.2; *p*<0.0001) as vehicle-treated animals performed poorly in this test burying significantly less marbles than the WT animals, like has been previously described (Segura-Puimedon et al., 2014). We observed a significant effect of treatment (F_3,66_ = 4.038; *p* = 0.0107) mainly due to a increased number of buried marbles in VERCUR treatment regardless of genotype. Treated CD animals showed altered behavior after any treatments, since we did not observe a significant interaction between genotype and treatment (F_3,66_ = 0.1146; *p* = 0.9512). (Figure 3).

**Figure 3.**
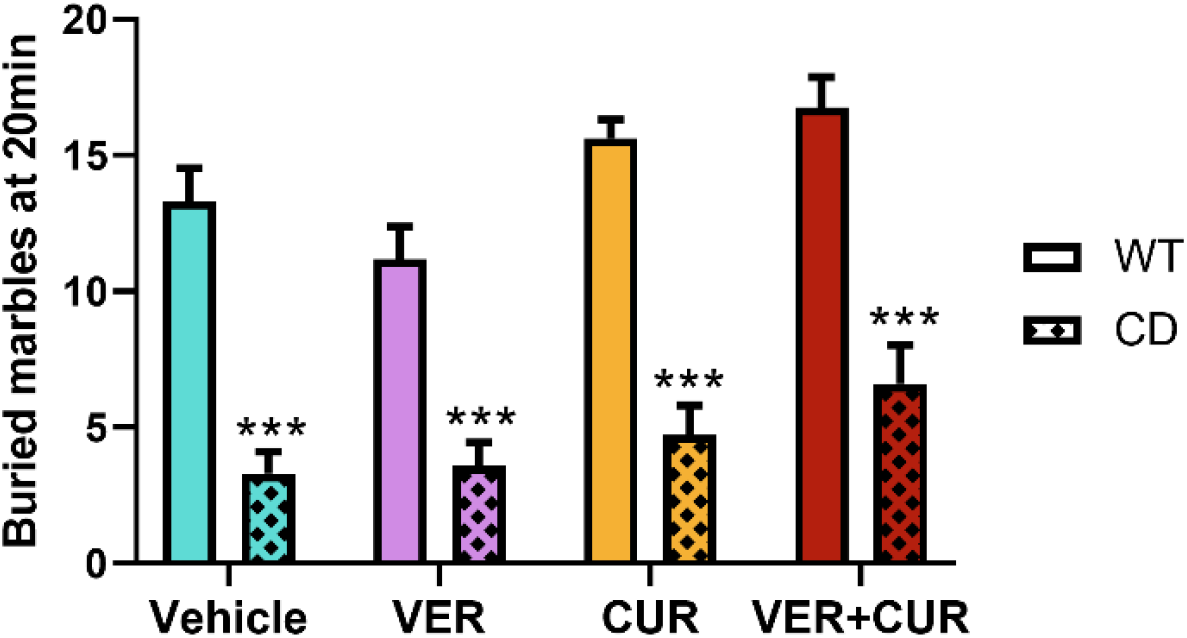
Anxiety-like behavior presented by CD animals is not affected by treatment with verapamil and curcumin. The number of buried marbles at the end point of the marble-burying test was significantly lower in CD animals compared to WT animals (effect of genotype F_1,66_=144.9; *p*<0.0001), and this effect was not prevented by any of the treatments (effect of treatment F_3,66_=5.604; *p*=0.0017). Data are presented as mean ± SEM of n=7-11 mice. *p* values are shown with asterisks indicating values that are significantly different in individual group comparisons (two-way ANOVA followed by Dunnett’s post hoc-test). \*\*\**p*<0.001.

### 3.4 Neuroanatomical features of CD mice do not change after VERCUR cotreatment

CD animals present a reduced brain weight, a disorganization in the anatomical of cortex neuronal layers, a reduction in the number of YFP+ expressing neurons and a decreased spine density (Segura-Puimedon et al., 2014; Borralleras et al., 2015; Ortiz-Romero et al., 2018). The aforementioned positive effects of VERCUR cotreatment in behavior led us to analyze the neuroanatomical characteristics of cotreated CD mice to evaluate if this was correlated with any changes at a neuroanatomical level.

In one hand, we observed that none of treatments had any effect in the recovery of a normal brain weight with a significant effect of genotype (F_1,54_=139.8, *p*<0.0001) but no effect of treatment (F_1,54_=1.606, *p*=0.2105) (Figure 4A, Supplemental Figure 5A). In the other, CD animals presented a significant reduction of the number of YFP+ neurons in both the motor cortex (effect of genotype F_1,25_=168.5, *p*<0.0001) and the hippocampus (effect of genotype F_1,25_=218.8, *p*<0.0001). Any of the treatments had any effect on the recovery of the number of YFP+ neurons (effect of treatment in motor cortex: F_3,25_=2.681, *p*=0.0686; effect of treatment in hippocampus: F_3,25_=1.426, *p*=0.2586) (Figure 4B, Supplemental Figure 5B and C). Moreover, the analysis VERCUR-cotreated animals revealed no differences in spine density and length showed neither in motor cortex nor in hippocampus (Figure 4C and D).

**Figure 4.**
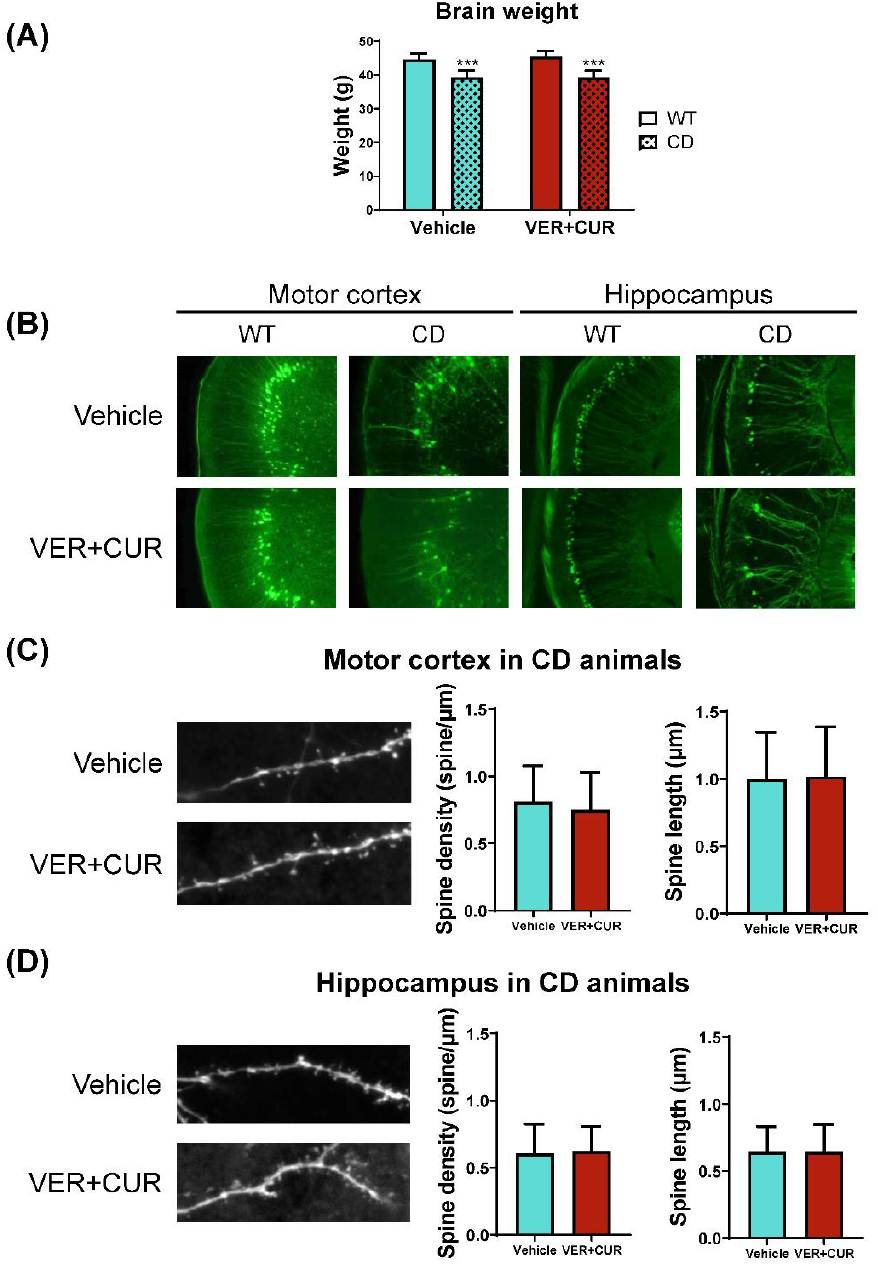
Treatment with verapamil and curcumin does not affect neuroanatomical features of CD mice. **(A)** Brain weight of VER+CUR treated CD animals was significantly reduced when compared to WT mice. A two-way ANOVA indicated a significant effect of genotype (F_1,54_=139.8, *p*<0.0001) but no effect of treatment (F_1,54_=1.606, *p*=0.2105). Data are presented as mean ± SEM of n=13-15 mice. *p* values are shown with asterisks indicating values that are significantly different in individual group comparisons (two-way ANOVA followed by Tukey’s*post hoc* test). ***p<0.001. **(B)** Representative images of motor cortex and hippocampus in vehicle- and VER+CUR-treated WT and CD animals. **(C, D)** Representative images and quantification of spine density and spine length in the motor cortex **(C)** and hippocampus **(D)** of vehicle- and VER+CUR treated CD animals. Data are presented as mean ± SEM of n=3-4 mice. Comparison with Mann-Whitney U-test showed no significant differences.

### 3.5 VERCUR treatment prevents the increased microglial activation presented by CD animals

The evaluation of the neuroinflammatory state of the brain was assessed by measuring the amount of activated microglia in different brain regions. CD animals showed an increase in the density of IBA1-positive cells, which is normalized after cotreatment with verapamil and curcumin. These results were observed in the motor cortex (effect of treatment F_1,13_=20.67; *p*=0.0005) (Figure 5A), the somatosensory cortex (effect of treatment F_1,13_=28.84; *p*=0.0001) (Figure 5B) and the hippocampus (effect of treatment F_1,13_=15.26; *p*=0.0018) (Figure 5C).

**Figure 5.**
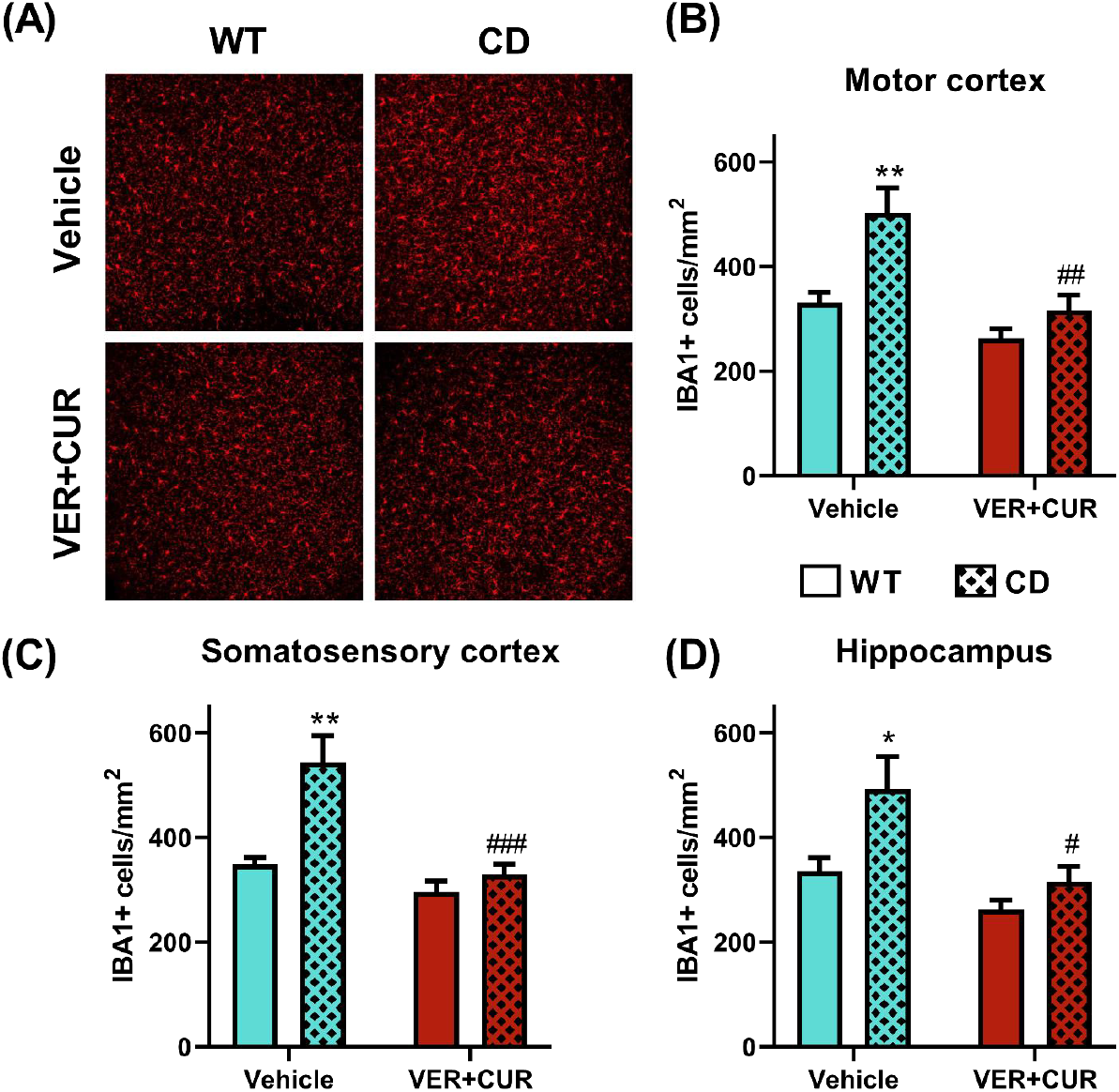
Treatment with verapamil and curcumin prevents the increased microglial activation presented by CD animals. **(A)** Representative images of IBA1 immunostaining in motor cortex from vehicle- and VER+CUR-treated WT and CD animals. **(B, C, D)** Quantification of IBA1 positive cells showed a significant effect of VER+CUR treatment in all the analyzed regions, which included **(B)** the motor cortex (effect of treatment F_1,13_=20.67; *p*=0.0005), **(C)** the somatosensory cortex (effect of treatment F_1,13_=28.84; *p*=0.0001) and **(D)** the hippocampus (effect of treatment F_1,13_=15.26; *p*=0.0018). Data are presented as mean ± SEM of n=3-5 mice. *P* values are shown with asterisks (effect of genotype) and hashes (effect of treatment) indicating values that are significantly different in individual group comparisons (two-way ANOVA followed by Dunnett’s *post hoc* test. *,#*p*<0.05, **,##*p*<0.01, ***,###*p*<0.001.

### 3.6 Combinatorial treatment recovers altered gene expression in cortex of CD animals

To identify the molecular causes that could be related to the improvement in certain aspects of the behavioral phenotype in VERCUR-treated animals, we investigated differentially expressed genes (DEG) by performing an RNA-seq analysis of cerebral cortex from VERCUR-treated CD mice *versus* vehicle-treated CD and WT mice.

First, we conducted a targeted analysis of the genes in the WSCR locus. Of the 26 genes that make up the WSCR, only 19 were measurably expressed in the adult mouse cortex. As expected,, all genes in the WSCR region showed a decrease in RNA amount in CD animals (Supplemental figure 2). We found 737 genes to be deregulated. The magnitude of these changes was generally small, with a fold-change ranking from 0,35 to 3,72. Differential expression analysis of vehicle-treated CD and WT samples identified 424 genes with significantly increased expression and 313 genes with a decreased one. The expression of the top 100 DEG clearly identified the two genotypes (Figure 6, Supplemental table 3). When ordered by statistical significance, up- and down-regulated genes had similar *p*-values.

**Figure 6.**
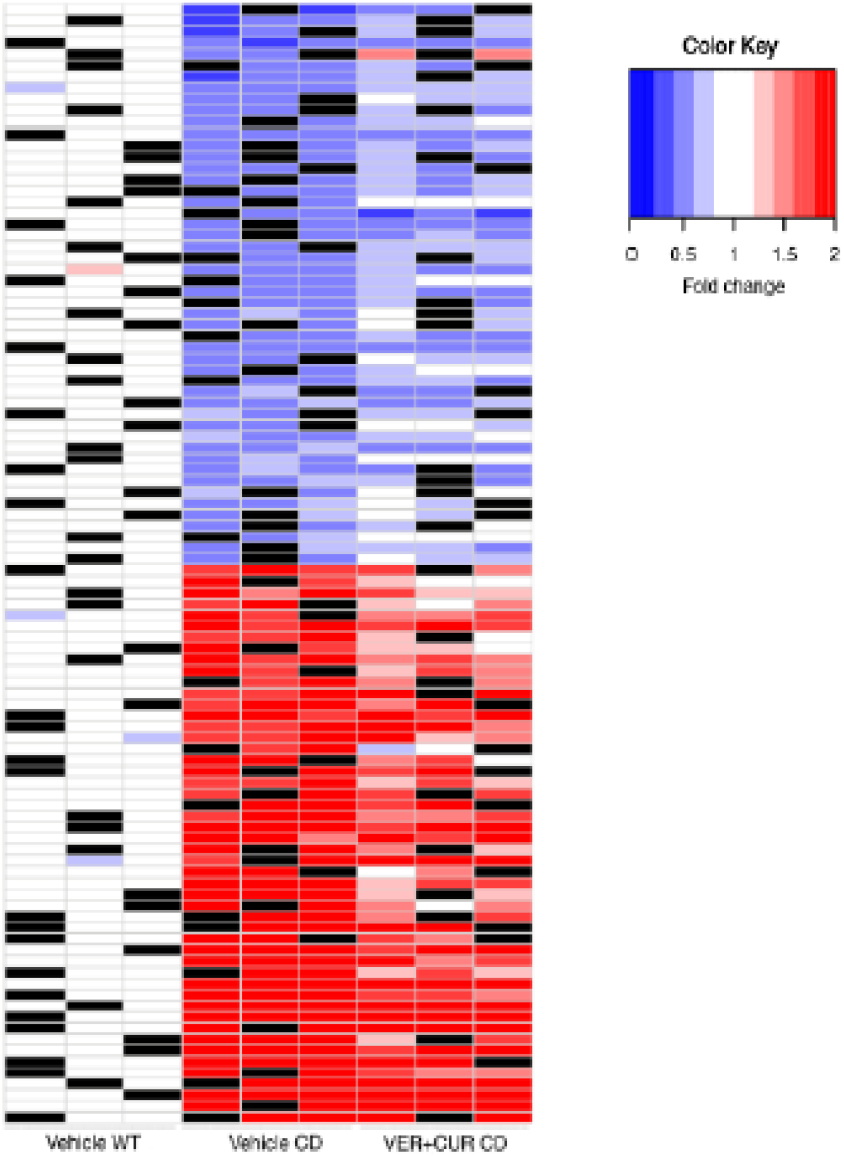
VERCUR treatament recovers altered gene expression in cortex of CD mice. Heatmap of the top 100 differentially expressed genes in cortex. Figure shows the to 50 up- and down-regulated genes with the exclusion of genes included in the WBSCR. Black: no sample.

In order to investigate functional associations of the DEG, we performed a gene set enrichment analysis (GSEA) using the CPDB tool to identify significantly differentiated KEGG/Reactome pathways. First, the enrichment analysis was performed between vehicle-treated WT and CD mice. Results showed 15 pathways that were significantly altered (*q*≤0.05) in CD compared to WT mice (Supplemental Table 4). Among them, the highest enrichment (*q*<0.015) was observed in pathways related to G protein coupled receptor (GPCR) ligand binding, extracellular matrix organization and mitogen-activated protein kinase (MAPK) signaling.

A second analysis was carried out to investigate the correlation between phenotypic and molecular rescue caused by the cotreatment. We performed a second analysis comparing, among the DEG between vehicle-treated groups, those that were not significantly different between vehicle-treated WT and VERCUR-treated CD. The expression of the 35.7% of the genes was recovered by the treatment (Supplemental table 5). Results of the GSEA analysis with these genes showed 4 differentiated pathways consistent with treatment rescue (*q* ≤ 0.05) (Table 1), and MAPK signaling related pathways were found among them (*q*=0.0138).

**Table 1.**
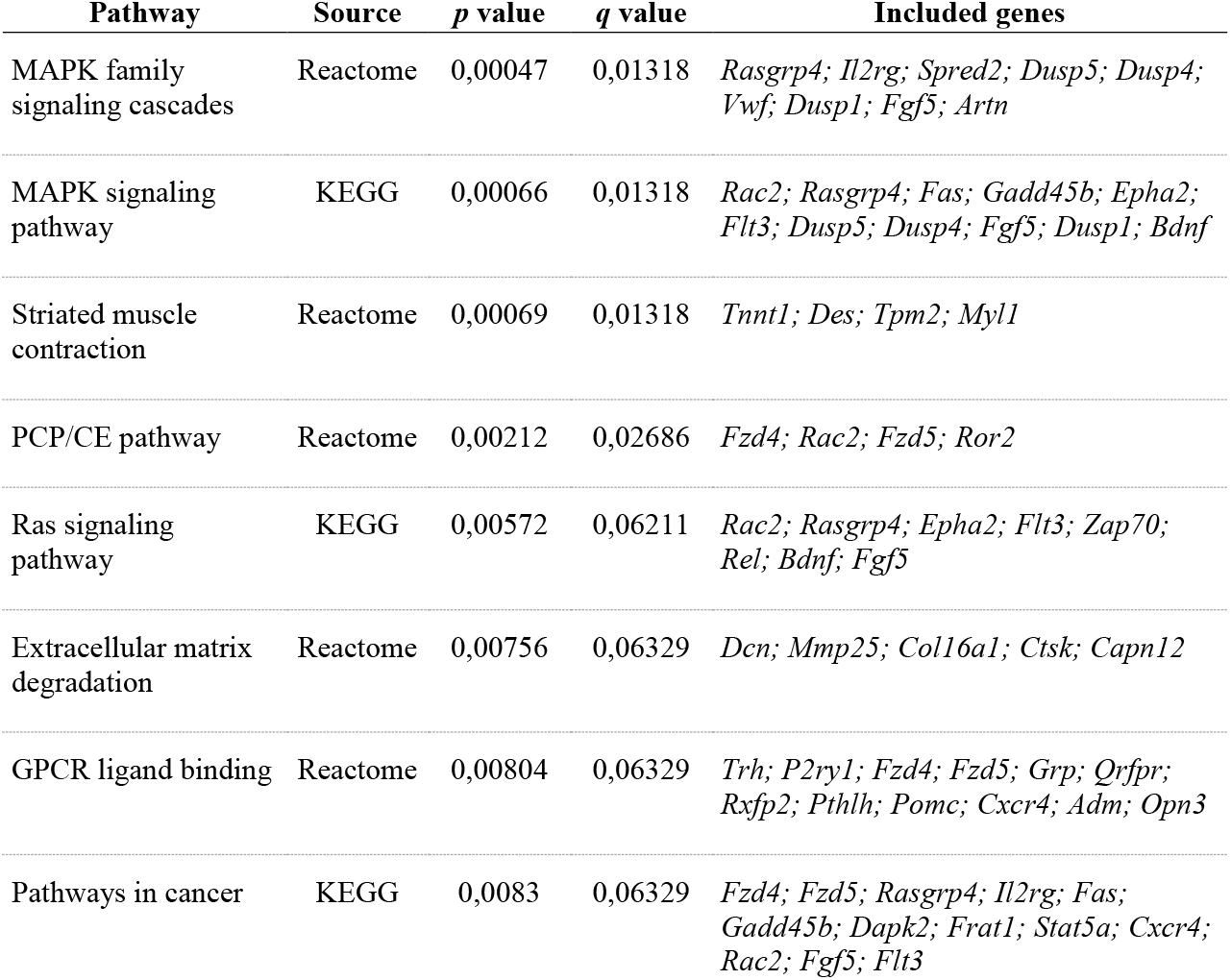
Pathway involvement of the differentially expressed genes recovered after treatment

## 4 Discussion

In this work we analyzed the effect of a treatment combining curcumin, the most abundant phenol in turmeric, and verapamil, a widely used medication, on the CD murine model of WBS. Neurobehavioral phenotype of CD mice recapitulates most of the cognitive features observed in WBS patients, such as motor discoordination, anxiety-like behavior and hypersociability (Segura-Puimedon et al., 2014).

The most significant improvement after treatment was observed in motor discoordination, evaluated with the rotarod test. Due to its effect in the modulation of calcium channels, verapamil has been studied as a therapeutic approach for different motor impairments such as dystonia or alcohol-induced motor discoordination. However, it has not given positive results in any of the cases (Jinnah et al., 2000; Baliño et al., 2010). In the case of curcumin, results have been contradictory. On the one hand, it has been described to significantly improve gait impairments in a model of α-synuclein deficit (Spinelli et al., 2015). On the other hand, it has been associated to detrimental effects in motor coordination in a Huntington’s disease murine model, without affecting locomotor function or muscle strength (Hickey et al., 2012). In our study, none of the individual treatments had any effect, while the combinatorial treatment prevented motor discoordination in CD animals. This result reinforces the idea that the mechanisms of action of both molecules are different and complementarity is needed to obtain positive effects on the CD phenotype.

Previous studies have defined that curcumin causes an increase in the exploratory behavior of stressed rats in an open field test (Vasileva et al., 2018). Our results in the evaluation of sociability agree with this finding, as we saw an increased exploration time of the empty container in the CUR-receiving groups, either in single administration or in combination with VER. This effect was observed both in WT and CD animals. However, the exploration time of the occupied container was not proportionally increased in CUR-consuming CD mice, suggesting a loss of the interest in the intruder mouse and thus a positive effect of this molecule on the social behavior of CD animals.

Performance of treated CD animals in the MBT was slightly different from untreated controls, what may be an indicative of reduced anxiety-like / compulsive behavior. Nevertheless, this improvement was mild and CD mice did not reach a normal performance as observed in WT animals. Studies focusing on the relevance of verapamil for neurological phenotypes have described it to have anxiolytic effects in mouse and rat models (Gopala Krishna et al., 2001; Biala and Budzynska, 2006), probably by interacting with receptors densely distributed in the cortex, the amygdala and the hippocampus (Matsumoto et al., 1994). Studies using curcumin for the treatment of anxiety-related behaviors have obtained, again, contradictory results. While some authors suggest it has anti-anxiety properties (Gilhotra and Dhingra, 2010), others have not seen these effects (P. Kulkarni et al., 2011). In our study, neither of the single treatments caused any changes on CD mice performance. However, it is interesting to note that all studies addressing anxiolytic properties of this compounds have used the elevated plus maze test. Therefore, our use of the MBT may be evaluating a different type of anxiety or compulsive-like behavior (de Brouwer et al., 2019), and this tendency to improvement of double-treated CD animals in this test should be deeply explored.

Using RNA-seq we have identified a large number of differentially expressed genes (DEG) in cortical tissue between WT and CD animals, revealing that the hemizygous loss of WBSCR genes has widespread effects. An enrichment analysis performed with these DEG indicates that the changes observed in gene expression may impact pathways related to MAPK signaling, extracellular matrix conformation and GPCR ligand binding signaling. Aberrant signaling in all these pathways may have negative consequences in the neurocognitive phenotype.

MAPK signaling-related alterations are linked to downregulation of *Bdnf* expression. Activation of TrkB receptor by BDNF promotes downstream intracellular signaling cascades involving, among others, Ras/MAPK, PLC-γ and PI3K/Akt pathways. These pathways are involved in regulating a variety of processes including neural differentiation and maturation and synaptic function (Numakawa et al., 2018), and alterations in MAPK signaling pathways have been described to cause behavioral impairments (Reinecke et al., 2013). Thus, we suggest that the downregulation of *Bdnf* and its relation with MAPK signaling pathways may be at least partly responsible for the neurocognitive phenotype of CD animals, which is consistent with what has been described in previous studies (Segura-Puimedon et al., 2014; Borralleras et al., 2015; Ortiz-Romero et al., 2018).

Combinatorial VERCUR treatment caused a recovery of MAPK signaling-related pathways. We attributed this result to a combination of the different properties of each molecule. Curcumin has been described to induce BDNF expression (Zhang et al., 2012; Nam et al., 2014), thus suggesting that this molecule is the main responsible of the *Bdnf* increased expression in treated CD animals. Moreover, a BDNF increase in the motor related cortex has been associated with improvements in motor coordination in mouse (Inoue et al., 2018). However, previous studies in our group had determined that solely the increase of *Bdnf* expression is not enough to improve the cognitive features of these animals (Ortiz-Romero et al., 2018). It has been described that BDNF action is highly related to a tightly controlled ionic homeostasis (Li et al., 1999; Numakawa et al., 2001; Adasme et al., 2011). Ionic signaling, especially regarding calcium, is closely related to MAPK signaling pathways as they include many calcium-dependent proteins. This suggests that verapamil is also modifying these affected pathways by contributing to the regulation of ionic homeostasis, which is crucial for the proper action of BDNF.

Interestingly, the effects of the VERCUR treatment over the behavioral phenotype of CD mice are similar to the effects of a genetic rescue of *Gtf2i* in these animals (Borralleras et al., 2015). In that study, the increase of *Gtf2i* expression levels by gene therapy correlated with a recovery of *Bdnf* levels in hippocampus. *Gtf2i* function has been tightly related to the control of PI3K/Akt pathway (Segura-Puimedon et al., 2013). All this reinforces the idea that BDNF deficiency and alterations in the downstream pathways could have a main role in the neurocognitive impairments of CD animals, and that their recovery is fundamental for the design of a good therapeutic approach.

GPCR ligand binding and extracellular matrix conformation related pathways were also found altered in cortical tissue, which may be contributing to the cognitive phenotype presented by CD animals.

GPCR activation is involved in a wide variety of physiological processes that include behavioral modulation through neurotransmitters-dependent activation, and deregulation of this process has been related with mood disorders and depression (Grammatopoulos, 2017). Conservation of extracellular matrix structures in the central nervous system has been described as fundamental for processes like cell migration, neural growth, synaptogenesis and synaptic plasticity, among others (Song and Dityatev, 2018). Interestingly, routes regarding extracellular matrix function have already been described as disease-relevant pathways in transcriptomic analysis performed on human WBS cell lines (Henrichsen et al., 2011; Khattak et al., 2015). Nevertheless, treatment was not able to fully recover the deficits found in these pathways.

Thus, results suggest that the effect of treatment on MAPK signaling-related pathways is contributing to the recovery of the functionality of synaptic signaling, which may be correlated with the improvements seen in behavior. However, the partial effects of treatment on the recovery of other altered pathways like GPCR ligand binding and extracellular matrix conformation may be correlated with the lack of improvement of the neuroanatomical features observed in the evaluation of brain morphology. We saw a reduction of the number of YFP+ expressing neurons in CD animals, as had already been reported (Segura-Puimedon et al., 2014). Similar reductions have been observed in other neurodevelopmental diseases mouse models (Stuss et al., 2012), although it has not been clarified if the reduced number of YFP+ neurons reflects an altered proportion of pyramidal neurons subclasses or is a consequence of aberrant activation of the transgene. Treatment had no effect on the recovery of a normal number of YFP+ neurons, and the study of dendritic spines in those cells also showed an absence of effect on the length or density of synaptic spines in CD mice.

Moreover, we also observed an increase in the amount of activated microglia in different brain regions of CD animals, which was prevented by treatment. Interestingly, alterations in activated glia density have also been observed in other mice models for different neurodevelopmental diseases (Lee et al., 2019; Matta et al., 2020). Curcumin has been previously related with the increase of *Nrf2* expression and nuclear translocation (Tapia et al., 2012; Fattori et al., 2015), suggesting that this endogenous anti-inflammatory pathway is modulating this response in CD animals, which correlates with similar findings in this model (Ortiz-Romero et al., 2018).

Altogether, the evaluation of gene expression in cerebral cortex contributes to the unraveling of mechanisms involved in the WBS cognitive profile. Nonetheless, it has to be taken into account that the same deletion is present in all tissues of the organism. Thus, it has to be considered that the observed alterations may have consequences in other organs and systems besides the brain, as all of the affected pathways are ubiquitously expressed and have a high relevance in a wide variety of cell processes.

In conclusion, we suggest that the hemizygous loss of WBSCR in cerebral cortex of CD mice has a direct effect on the neuroinflammatory state of the brain, as well as in the expression of some genes related to synaptic signaling or extracellular matrix structure, that are crucial for a proper neural function. This may be at least partly responsible for the behavioral phenotype observed in CD animals. A treatment combining verapamil and curcumin is able to address different molecular targets and rescue some of those pathways, being a promising therapeutic approach for the cognitive phenotype of WBS patients.

## Supporting information

Supplemental Ortiz-Romero 2021

## 5 Conflict of Interest

The authors declare that the research was conducted in the absence of any commercial or financial relationships that could be construed as a potential conflict of interest.

## 6 Author Contributions

Substantial contributions to the conception/ design of the work:GE-LPJ-VC

The acquisition, analysis, and interpretation of data for the work: POR-VC

Drafting the work : POR-VC

Revising the work critically for important intellectual content: GE-LPJ

Final approval of the version to be published; POR-GE-LPJ-VC

Agreement to be accountable for all aspects of the work in ensuring that questions related to the accuracy or integrity of any part of the work are appropriately investigated and resolved:POR-GE-LPJ-VC

## 7 Funding

This work was supported by Ministerio de Ciencia e Innovación [SAF2017-83039-R to GE] and [SAF2016-78508-R (AEI/MINEICO/FEDER, UE) to VC]; from Association”Autour des Williams” to VC; and Generalitat de Catalunya [2017-SGR1794] to VC

## 8 Acknowledgments

We thanks Marta Linares for Technical assistance.

## Notes

### Competing Interest Statement

The authors have declared no competing interest.

## References

Adasme, T., Haeger, P., Paula-Lima, A. C., Espinoza, I., Casas-Alarcón, M. M., Carrasco, M. A., et al. (2011). Involvement of ryanodine receptors in neurotrophin-induced hippocampal synaptic plasticity and spatial memory formation. Proc. Natl. Acad. Sci. U. S. A. 108, 3029–3034. doi:10.1073/pnas.1013580108.

Aubry, A. V., Khandaker, H., Ravenelle, R., Grunfeld, I. S., Bonnefil, V., Chan, K. L., et al. (2019). A diet enriched with curcumin promotes resilience to chronic social defeat stress. Neuropsychopharmacology 44, 733–742. doi:10.1038/s41386-018-0295-2.

Babu, A., Prasanth, K. G., and Balaji, B. (2015). Effect of curcumin in mice model of vincristine-induced neuropathy. Pharm. Biol. 53, 838–848. doi:10.3109/13880209.2014.943247.

Baliño, P., Pastor, R., and Aragon, C. M. G. (2010). Participation of L-type calcium channels in ethanol-induced behavioral stimulation and motor incoordination: Effects of diltiazem and verapamil. Behav. Brain Res. 209, 196–204. doi:10.1016/j.bbr.2010.01.036.

Bayés, M., Magano, L. F., Rivera, N., Flores, R., and Pérez Jurado, L. A. (2003). Mutational mechanisms of williams-beuren syndrome deletions. Am. J. Hum. Genet. 73, 131–151. doi:10.1086/376565.

Bellamy, W. T. (1996). P-glycoproteins and multidrug resistance. Annu. Rev. Pharmacol. Toxicol. 36, 161–183. doi:10.1146/annurev.pharmtox.36.1.161.

Bellugi, U., Lichtenberger, L., Jones, W., Lai, Z., and St. George, M. (2000). I. The neurocognitive profile of Williams syndrome: A complex pattern of strengths and weaknesses. J. Cogn. Neurosci. 12, 7–29. doi:10.1162/089892900561959.

Biala, G., and Budzynska, B. (2006). Effects of acute and chronic nicotine on elevated plus maze in mice: Involvement of calcium channels. Life Sci. 79, 81–88. doi:10.1016/j.lfs.2005.12.043.

Biala, G., and Kruk, M. (2008). Calcium channel antagonists suppress cross-tolerance to the anxiogenic effects of d-amphetamine and nicotine in the mouse elevated plus maze test. Prog. Neuro-Psychopharmacology Biol. Psychiatry 32, 54–61. doi:10.1016/j.pnpbp.2007.07.006.

Borralleras, C., Sahun, I., Pérez-Jurado, L. A., and Campuzano, V. (2015). Intracisternal Gtf2i gene therapy ameliorates deficits in cognition and synaptic plasticity of a mouse model of williams-beuren syndrome. Mol. Ther. 23. doi:10.1038/mt.2015.130.

de Brouwer, G., Fick, A., Harvey, B. H., and Wolmarans, D. W. (2019). A critical inquiry into marble-burying as a preclinical screening paradigm of relevance for anxiety and obsessive-compulsive disorder: Mapping the way forward. Cogn. Affect. Behav. Neurosci. 19, 1–39. doi:10.3758/s13415-018-00653-4.

Doyle, T. F., Bellugi, U., Korenberg, J. R., and Graham, J. (2004). “Everybody in the World Is My Friend” Hypersociability in Young Children with Williams Syndrome. Am. J. Med. Genet. 124 A, 263–273. doi:10.1002/ajmg.a.20416.

Elliott, W. J., and Ram, C. V. S. (2011). Calcium channel blockers. J. Clin. Hypertens. 13, 687–689. doi:10.1111/j.1751-7176.2011.00513.x.

Fattori, V., Pinho-Ribeiro, F. A., Borghi, S. M., Alves-Filho, J. C., Cunha, T. M., Cunha, F. Q., et al. (2015). Curcumin inhibits superoxide anion-induced pain-like behavior and leukocyte recruitment by increasing Nrf2 expression and reducing NF-κB activation. Inflamm. Res. 64, 993–1003. doi:10.1007/s00011-015-0885-y.

Feng, G., Mellor, R. H., Bernstein, M., Keller-Peck, C., Nguyen, Q. T., Wallace, M., et al. (2000). Imaging Neuronal Subsets in Transgenic Mice Expressing Multiple Spectral Variants of GFP variants with altered spectral properties and improved translational efficiency, thermostability, and quantum yield. As a result of these favorabl. Neuron 28, 41–51.

Franco-Robles, E., Campos-Cervantes, A., Murillo-Ortiz, B. O., Segovia, J., López-Briones, S., Vergara, P., et al. (2014). Effects of curcumin on brain-derived neurotrophic factor levels and oxidative damage in obesity and diabetes. Appl. Physiol. Nutr. Metab. 39, 211–218. doi:10.1139/apnm-2013-0133.

Gilhotra, N., and Dhingra, D. (2010). GABAergic and nitriergic modulation by curcumin for its antianxiety-like activity in mice. Brain Res. 1352, 167–175. doi:10.1016/j.brainres.2010.07.007.

Gopala Krishna, H. N., Kumar, K. B., and Karanth, K. S. (2001). The anxiolytic activity of calcium channel antagonists in experimental models of anxiety in rats. Indian J. Pharmacol. 33, 267–271.

Grammatopoulos, D. K. (2017). Regulation of G-protein coupled receptor signalling underpinning neurobiology of mood disorders and depression. Mol. Cell. Endocrinol. 449, 82–89. doi:10.1016/j.mce.2017.02.013.

Green, T., Avda, S., Dotan, I., Zarchi, O., Basel-Vanagaite, L., Zalsman, G., et al. (2012). Phenotypic psychiatric characterization of children with Williams syndrome and response of those with ADHD to methylphenidate treatment. Am. J. Med. Genet. Part B Neuropsychiatr. Genet. 159 B, 13–20. doi:10.1002/ajmg.b.31247.

Hay, E., Lucariello, A., Contieri, M., Esposito, T., De Luca, A., Guerra, G., et al. (2019). Therapeutic effects of turmeric in several diseases: An overview. Chem. Biol. Interact. 310. doi:10.1016/j.cbi.2019.108729.

Henrichsen, C. N., Csárdi, G., Zabot, M. T., Fusco, C., Bergmann, S., Merla, G., et al. (2011). Using transcription modules to identify expression clusters perturbed in Williams-Beuren Syndrome. PLoS Comput. Biol. 7. doi:10.1371/journal.pcbi.1001054.

Hickey, M. A., Zhu, C., Medvedeva, V., Lerner, R. P., Patassini, S., Franich, N. R., et al. (2012). Improvement of neuropathology and transcriptional deficits in CAG 140 knock-in mice supports a beneficial effect of dietary curcumin in Huntington’s disease. Mol. Neurodegener. 7, 12. doi:10.1186/1750-1326-7-12.

Inoue, T., Ninuma, S., Hayashi, M., Okuda, A., Asaka, T., and Maejima, H. (2018). Effects of longterm exercise and low-level inhibition of GABAergic synapses on motor control and the expression of BDNF in the motor related cortex. Neurol. Res. 40, 18–25. doi:10.1080/01616412.2017.1382801.

Jin, M., Park, S. Y., Shen, Q., Lai, Y., Ou, X., Mao, Z., et al. (2018). Anti-neuroinflammatory effect of curcumin on Pam3CSK4-stimulated microglial cells. Int. J. Mol. Med. 41, 521–530. doi:10.3892/ijmm.2017.3217.

Jinnah, H. A., Sepkuty, J. P., Ho, T., Yitta, S., Drew, T., Rothstein, J. D., et al. (2000). Calcium channel agonists and dystonia in the mouse. Mov. Disord. 15, 542–551. doi:10.1002/1531-8257(200005)15:3<542::AID-MDS1019>3.0.CO;2-2.

Khattak, S., Brimble, E., Zhang, W., Zaslavsky, K., Strong, E., Ross, P. J., et al. (2015). Human induced pluripotent stem cell derived neurons as a model for Williams-Beuren syndrome. Mol. Brain 8, 1–11. doi:10.1186/s13041-015-0168-0.

Ko, E. A., Park, W. S., Son, Y. K., Ko, J. H., Choi, T. H., Jung, I. D., et al. (2010). Calcium channel inhibitor, verapamil, inhibits the voltage-dependent K + channels in rabbit coronary smooth muscle cells. Biol. Pharm. Bull. 33, 47–52. doi:10.1248/bpb.33.47.

Kopp, N., McCullough, K., Maloney, S. E., and Dougherty, J. D. (2019). Gtf2i and Gtf2ird1 mutation do not account for the full phenotypic effect of the Williams syndrome critical region in mouse models. Hum. Mol. Genet. 28, 3443–3465. doi:10.1093/hmg/ddz176.

Lee, F. H. F., Lai, T. K. Y., Su, P., and Liu, F. (2019). Altered cortical Cytoarchitecture in the Fmr1 knockout mouse. Mol. Brain 12, 1–12. doi:10.1186/s13041-019-0478-8.

Li, H. H., Roy, M., Kuscuoglu, U., Spencer, C. M., Halm, B., Harrison, K. C., et al. (2009). Induced chromosome deletions cause hypersociability and other features of Williams-Beuren syndrome in mice. EMBO Mol. Med. 1, 50–65. doi:10.1002/emmm.200900003.

Li, H. S., Xu, X. Z. S., and Montell, C. (1999). Activation of a trpc3-dependent cation current through the neurotrophin bdnf. Neuron 24, 261–273. doi:10.1016/S0896-6273(00)80838-7.

Martens, M. A., Seyfer, D. L., Andridge, R. R., Foster, J. E. A., Chowdhury, M., McClure, K. E., et al. (2012). Parent report of antidepressant, anxiolytic, and antipsychotic medication use in individuals with Williams syndrome: Effectiveness and adverse effects. Res. Dev. Disabil. 33, 2106–2121. doi:10.1016/j.ridd.2012.06.006.

Martens, M. A., Wilson, S. J., and Reutens, D. C. (2008). Research Review: Williams syndrome: A critical review of the cognitive, behavioral, and neuroanatomical phenotype. J. Child Psychol. Psychiatry Allied Discip. 49, 576–608. doi:10.1111/j.1469-7610.2008.01887.x.

Matsumoto, Y., Kataoka, Y., and Miyazaki, A. (1994). ejp. 264, 107–110.

Matta, S. M., Moore, Z., Walker, F. R., Hill-Yardin, E. L., and Crack, P. J. (2020). An altered glial phenotype in the NL3R451C mouse model of autism. Sci. Rep. 10, 1–13. doi:10.1038/s41598-020-71171-y.

Nam, S. M., Choi, J. H., Yoo, D. Y., Kim, W., Jung, H. Y., Kim, J. W., et al. (2014). Effects of curcumin (Curcuma longa) on learning and spatial memory as well as cell proliferation and neuroblast differentiation in adult and aged mice by upregulating brain-derived neurotrophic factor and CREB signaling. J. Med. Food 17, 641–649. doi:10.1089/jmf.2013.2965.

Numakawa, T., Matsumoto, T., Adachi, N., Yokomaku, D., Kojima, M., Takei, N., et al. (2001). Brain-derived neurotrophic factor triggers a rapid glutamate release through increase of intracellular Ca2+ and Na+ in cultured cerebellar neurons. J. Neurosci. Res. 66, 96–108. doi:10.1002/jnr.1201.

Numakawa, T., Odaka, H., and Adachi, N. (2018). Actions of brain-derived neurotrophin factor in the neurogenesis and neuronal function, and its involvement in the pathophysiology of brain diseases. Int. J. Mol. Sci. 19. doi:10.3390/ijms19113650.

Ortiz-Romero, P., Borralleras, C., Bosch-Morató, M., Guivernau, B., Albericio, G., Muñoz, F. J., et al. (2018). Epigallocatechin-3-gallate improves cardiac hypertrophy and short-term memory deficits in a Williams-Beuren syndrome mouse model. PLoS One 13. doi:10.1371/journal.pone.0194476.

P. Kulkarni, A., A. Govender, D., J. Kotwal, G., and A. Kellaway, L. (2011). Modulation of Anxiety Behavior by Intranasally Administered Vaccinia Virus Complement Control Protein and Curcumin in a Mouse Model of Alzheimers Disease. Curr. Alzheimer Res. 8, 95–113. doi:10.2174/156720511794604598.

Quartermain, D., and Garcia DeSoria, V. (2001). The effects of calcium channel antagonists on short- and long-term retention in mice using spontaneous alternation behavior. Neurobiol. Learn. Mem. 76, 117–124. doi:10.1006/nlme.2000.3981.

Reinecke, K., Herdegen, T., Eminel, S., Aldenhoff, J. B., and Schiffelholz, T. (2013). Knockout of c-Jun N-terminal kinases 1, 2 or 3 isoforms induces behavioural changes. Behav. Brain Res. 245, 88–95. doi:10.1016/j.bbr.2013.02.013.

Sanei, M., and Saberi-Demneh, A. (2019). Effect of curcumin on memory impairment: A systematic review. Phytomedicine 52, 98–106. doi:10.1016/j.phymed.2018.06.016.

Segura-Puimedon, M., Borralleras, C., Pérez-Jurado, L. A., and Campuzano, V. (2013). TFII-I regulates target genes in the PI-3K and TGF-β signaling pathways through a novel DNA binding motif. Gene 527, 529–536. doi:10.1016/j.gene.2013.06.050.

Segura-Puimedon, M., Sahún, I., Velot, E., Dubus, P., Borralleras, C., Rodrigues, A. J., et al. (2014). Heterozygous deletion of the Williams-Beuren syndrome critical interval in mice recapitulates most features of the human disorder. Hum. Mol. Genet. 23, 6481–6494. doi:10.1093/hmg/ddu368.

Song, I., and Dityatev, A. (2018). Crosstalk between glia, extracellular matrix and neurons. Brain Res. Bull. 136, 101–108. doi:10.1016/j.brainresbull.2017.03.003.

Spinelli, K. J., Osterberg, V. R., Meshul, C. K., Soumyanath, A., and Unni, V. K. (2015). Curcumin treatment improves motor behavior in α-synuclein transgenic mice. PLoS One 10, 1–20. doi:10.1371/journal.pone.0128510.

Strømme, P., Bjømstad, P. G., and Ramstad, K. (2002). Prevalence estimation of Williams syndrome. J. Child Neurol. 17, 269–271. doi:10.1177/088307380201700406.

Stuss, D. P., Boyd, J. D., Levin, D. B., and Delaney, K. R. (2012). MeCP2 mutation results in compartment-specific reductions in dendritic branching and spine density in layer 5 motor cortical neurons of YFP-H mice. PLoS One 7, 1–11. doi:10.1371/journal.pone.0031896.

Tapia, E., Soto, V., Ortiz-Vega, K. M., Zarco-Márquez, G., Molina-Jijón, E., Cristóbal-García, M., et al. (2012). Curcumin induces Nrf2 nuclear translocation and prevents glomerular hypertension, hyperfiltration, oxidant stress, and the decrease in antioxidant enzymes in 5/6 nephrectomized rats. Oxid. Med. Cell. Longev. 2012. doi:10.1155/2012/269039.

Vasileva, L. V., Saracheva, K. E., Ivanovska, M. V., Petrova, A. P., Marchev, A. S., Georgiev, M. I., et al. (2018). Antidepressant-like effect of salidroside and curcumin on the immunoreactivity of rats subjected to a chronic mild stress model. FoodChem. Toxicol. 121, 604–611. doi:10.1016/j.fct.2018.09.065.

von Bohlen und Halbach, O., and von Bohlen und Halbach, V. (2018). BDNF effects on dendritic spine morphology and hippocampal function. Cell Tissue Res. 373, 729–741. doi:10.1007/s00441-017-2782-x.

Zhang, L., Xu, T., Wang, S., Yu, L., Liu, D., Zhan, R., et al. (2012). Curcumin produces antidepressant effects via activating MAPK/ERK-dependent brain-derived neurotrophic factor expression in the amygdala of mice. Behav. Brain Res. 235, 67–72. doi:10.1016/j.bbr.2012.07.019.

