## Supplemental Ortiz-Romero 2021 for "Verapamil/Curcumin treatment attenuates the behavioral alterations observed in Williams Syndrome mice by regulation of MAPK pathway and Microglia overexpression"

### *Supplementary Material*

#### Supplementary Figures

#### 1 Supplementary Figure 1

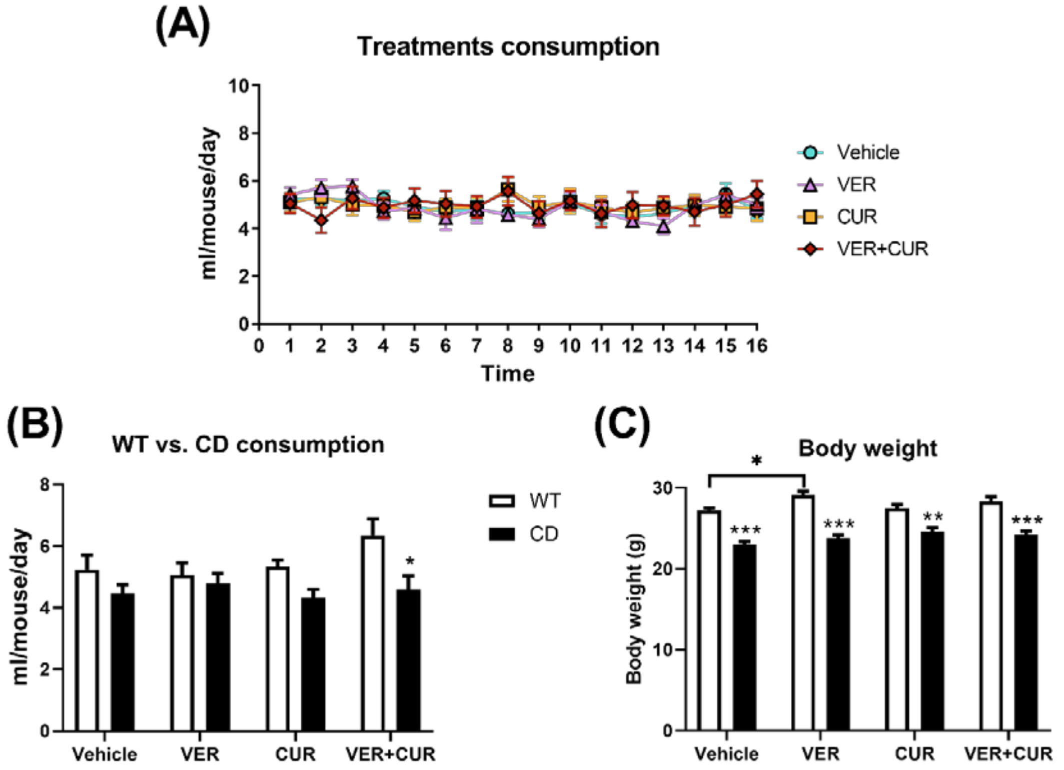


**Supplementary Figure 1. Treatment intake and body weight. (A)** Daily consumption (ml/mouse/day) of different treatments was equivalent (repeated measures ANOVA, no significant effect of treatment. $F_{2.005,30.08}$=0.4796, $p$=0.6242) and did not change over time (two-way ANOVA, no significant interaction between treatment and time. $F_{45,425}$=0.6025, $p$=0.9809). **(B)** Measurement of consumed amount of treatment (ml/mouse/day) showed a significant reduction in CD animals (two-way ANOVA, effect of genotype, $F_{1,41}$=11.11, $p$=0.0018). However, Bonferroni *post hoc* test revealed that the only significant difference was found between the VER+CUR-consuming WT and CD groups, and it was mainly due to an increase in the consumption of WT animals. **(C)** Body weights of CD mice were lower in comparison with WT mice (effect of genotype, $F_{1,84}$=146.4, $p$<0.0001) with a significant influence of treatment (effect of treatment, $F_{3,84}$=3.407, $p$=0.0213) due to a difference between vehicle- and VER-consuming WT animals. No differences were observed for the other groups of treatment. Data are presented as mean ± SEM of n=7-17 mice. *P* values are shown with asterisks indicating values that are significantly different in individual comparisons with Bonferroni or Tukey's *post hoc* tests. *$p$<0.05, **$p$<0.01, ***$p$<0.001.

### 2 Supplementary Figure 2

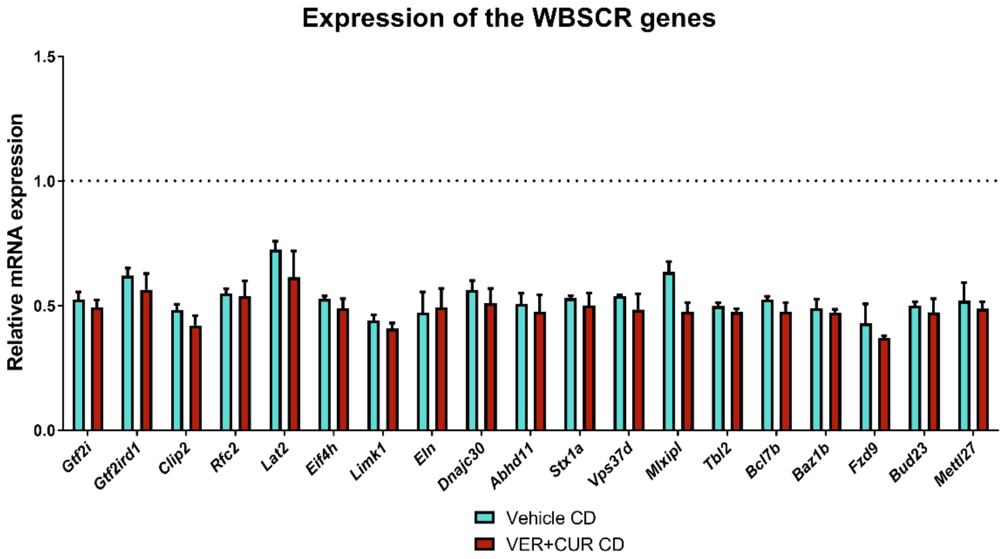


**Supplementary Figure 2. Expression of the WBS critical region genes.** RNA-seq expression analysis of genes contained in the WBSCR, analyzed from cerebral cortex, showed that the expression of all WBSCR genes was significantly downregulated in vehicle CD animals in comparison with the average expression in WT animals, represented in the graph with a dotted line (adjusted $p<0.05$ in all cases with Holm-Sidak correction after multiple t tests). Treatment had no effect in the recovery of normal expression levels for any of the genes as VER+CUR-CD expression levels were not different from vehicle-CD levels ($p>0.05$ in all cases with Holm-Sidak correction after multiple t tests). Data are represented as mean ± SEM of n=3 mice.

### 3 Supplementary Figure 3

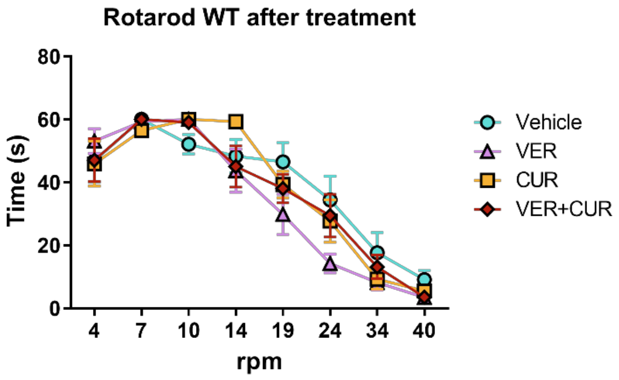


**Supplementary Figure 3. Rotarod results in WT treated groups.** Any of the treatments had any effect on the performance of WT animals in this test (two-way ANOVA, effect of treatment $F_{3,256}=1.918$, $p=0.1272$). Data are presented as mean ± SEM of n=7-11 mice.

### 4 Supplementary Figure 4

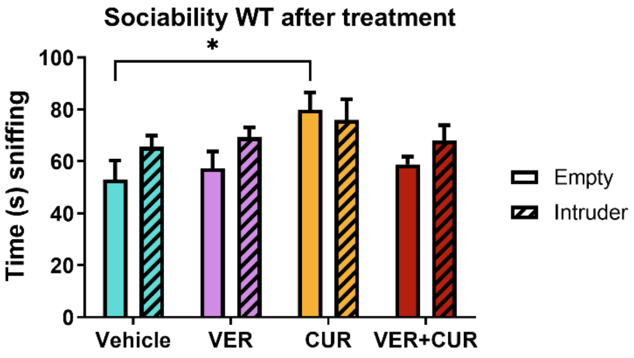


**Supplementary Figure 4. Sociability results in WT treated groups.** Evaluation of sociability in WT treated groups showed a significant effect of treatment (two-way ANOVA, effect of treatment $F_{3,66}=3.592$, $p=0.0181$) but mainly due to an increase in the exploration times of the CUR-treated group. Data are presented as mean ± SEM of n=7-11 mice. *P* values are shown with asterisks indicating values that are significantly different in individual comparisons with Tukey's *post hoc* test. *$p<0.05$.

### 5 Supplementary Figure 5

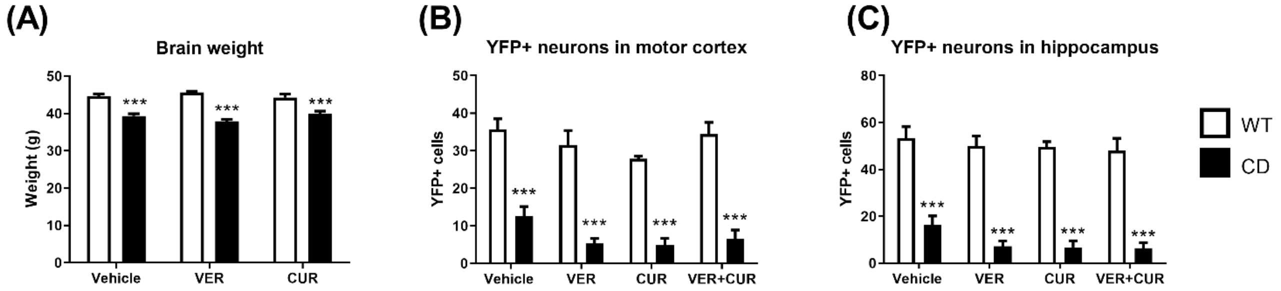


**Supplementary Figure 5. (A)** Brain weight evaluation showed that neither VER nor CUR single treatments had any effect on the recovery of a normal brain weight in CD animals. A two-way ANOVA indicated a significant effect of genotype ($F_{1,58}=136.9$, $p<0.0001$) but no effect of treatment ($F_{2,58}=0.1715$, $p=0.8428$). Data are presented as mean ± SEM of n=8-13 mice. *p* values are shown with asterisks indicating values that are significantly different in Bonferroni *post hoc* test. **(B,C)** Quantification of the number of YFP+ expressing neurons in **(B)** motor cortex and **(C)** hippocampus. CD animals presented a significant reduction of the number of YFP+ neurons in both the motor cortex (effect of genotype $F_{1,25}=168.5$, $p<0.0001$) and the hippocampus (effect of genotype $F_{1,25}=218.8$, $p<0.0001$). Any of the treatments had any effect on the recovery of the number of YFP+ neurons (effect of treatment in motor cortex: $F_{3,25}=2.681$, $p=0.0686$; effect of treatment in hippocampus: $F_{3,25}=1.426$, $p=0.2586$). Data are presented as mean ± SEM of n=3-5 mice. ***$p<0.001$.

### Supplementary Tables

**Supplementary Table 1. Primer sequences used for the expression analyses of *Vcam1* and *Nrf2***

| Gene | Sequence | Amplicon size | Location | Tm (ºC) |
|---|---|---|---|---|
| *Vcam1* | L: 5’-ATTTTCTGGGGCAGGAAGTT-3’ | 238bp | Exon 4 | 59.94 |
| | R: 5’- ACGTCAGAACAACCGAATCC-3’ | | Exon 5 | 59.97 |
| *Nrf2* (*Nfe212*) | L: 5’- CAGTCTTCACTGCCCCTCAT-3’ | 263bp | Exon 4 | 60.26 |
| | R: 5’- GTTGCCCACTTCTTTTTCCA-3’ | | Exon 5 | 60.09 |

**Supplementary Table 2. Summary of rotarod results.** Adjusted *p* values correspond to Tukey’s *posthoc* test after 2-way ANOVAS, one for each rpm value.

| | Veh WT vs Veh CD | Veh CD vs. VER+CUR CD | Veh WT vs. VER+CUR CD |
|---|---|---|---|
| **4rpm** | 0.2417 | >0.9999 | 0.2620 |
| **7rpm** | 0.0332 | 0.0332 | >0.9999 |
| **10rpm** | 0.0363 | 0.0254 | 0.9958 |
| **14rpm** | 0.0494 | 0.2227 | 0.9064 |
| **19rpm** | 0.0081 | 0.2421 | 0.4026 |
| **24rpm** | 0.2930 | 0.9808 | 0.4998 |
| **34rpm** | 0.0504 | 0.6488 | 0.4100 |
| **40rpm** | 0.2577 | 0.9747 | 0.4454 |

**Supplementary Table 3. Pathway involvement of the differentially expressed genes between vehicle-treated WT and CD mice.**

| Pathway | Source | *p* value | *q* value | Included genes |
|---|---|---|---|---|
| MAPK signaling pathway | KEGG | $6.66 \cdot 10^{-5}$ | 0.01009 | *Fas; Ntf3; Rac2; Rasgrp4; Hspb1; Fgf5; Gadd45a; Gadd45b; Nr4a1; Epha2; Vegfd; Dusp9; Il1r1; Relb; Dusp5; Dusp4; Bdnf; Dusp1; Flt3; Cacng6; Hspa1l* |
| GPCR ligand binding | Reactome | $8.55 \cdot 10^{-5}$ | 0.01009 | *Glp2r; Tbxa2r; Ccr1; Pf4; Ptgdr; Opn3; Cysltr1; Adm; Fzd4; Fzd5; Fzd9; Rxfp2; Pthlh; Cxcr4; Trh; Grp; Npffr1; Pomc; Grpr; P2ry1; Tacr3; Bdkrb2; Nts; Adrb3; Oprd1; Agtr2; Qrfpr; Gng8* |
| Extracellular matrix organization | Reactome | 0.00018 | 0.0144 | *Dcn; Ctsk; Adamts14; Col7a1; Itgb7; Tnxb; Capn12; Tmprss6; Mmp19; Mmp25; Adam12; Optc; Col28a1; Gdf5; Col6a1; Adam19; Col16a1; Col2a1; A2m; Col17a1* |
| Peptide ligand-binding receptors | Reactome | 0.0003 | 0.01494 | *Trh; Grpr; Bdkrb2; Grp; Qrfpr; Rxfp2; Npffr1; Ccr1; Pomc; Oprd1; Cxcr4; Agtr2; Tacr3; Nts; Pf4* |
| Class A/1 (Rhodopsin-like receptors) | Reactome | 0.00032 | 0.01494 | *Trh; Grpr; Bdkrb2; Grp; Tbxa2r; Tacr3; Qrfpr; Rxfp2; Npffr1; Ccr1; Pomc; Adrb3; Pf4; Oprd1; Cxcr4; Ptgdr; Agtr2; P2ry1; Opn3; Nts; Cysltr1* |
| Extracellular matrix degradation | Reactome | 0.00062 | 0.02108 | *Dcn; Ctsk; Mmp19; Tmprss6; Capn12; Optc; Mmp25; Col16a1; A2m; Col17a1* |
| Formation of fibrin clot (clotting cascade) | Reactome | 0.00063 | 0.02108 | *Pf4; F12; F13a1; Vwf; A2m; Serpind1* |
| NF-κβ signaling pathway | KEGG | 0.0011 | 0.03161 | *Lbp; Gadd45b; Nfkbia; Pidd1; Zap70; Relb; Card14; Lck; Il1r1* |
| Collagen chain trimerization | Reactome | 0.00121 | 0.03161 | *Col7a1; Col28a1; Col6a1; Col16a1; Col2a1; Col17a1* |
| Arachidonic acid metabolism | KEGG | 0.00164 | 0.03879 | *Fam213b; Pla2g2d; Gpx3; Pla2g3; Plb1; Hpgds; Alox12* |
| Striated muscle contraction | Reactome | 0.00251 | 0.05028 | *Tnnt1; Tpm2; Des; Tnnc1; Myl1* |
| Collagen biosynthesis and modifying enzymes | Reactome | 0.00256 | 0.05028 | *Col7a1; Col28a1; Col6a1; Col16a1; Col17a1; Col2a1; Adamts14* |
| Intrinsic pathway of fibrin clot formation | Reactome | 0.00277 | 0.05028 | *F12; Vwf; A2m; Serpind1* |
| RAF-independent MAPK1/3 activation | Reactome | 0.00328 | 0.05162 | *Dusp5; Dusp4; Dusp1; Dusp9* |

| Pathway | Source | *p* value | *q* value | Included genes |
|---|---|---|---|---|
| Histidine metabolism | KEGG | 0.00328 | 0.05162 | *Amdhd1; Aldh3b1; Cndp1; Aldh3b2* |
| PI3K-Akt signaling pathway | KEGG | 0.00413 | 0.06092 | *Ppp2r1b; Ntf3; Ddit4; Itgb7; Sgk1; Creb3l3; Nr4a1; Tnxb; Epha2; Flt3; Vegfd; Il2rg; Eif4ebp1; Col6a1; Vwf; Fgf5; Col2a1; Bdnf; Gng8* |
| Negative regulation of MAPK pathway | Reactome | 0.00455 | 0.06312 | *Dusp5; Dusp4; Ppp2r1b; Dusp1; Dusp9* |
| Pathways in cancer | KEGG | 0.00554 | 0.0727 | *Fas; Rasgrp4; Nfkbia; Flt3; Rara; Heyl; Fzd4; Zbtb16; Fzd9; Gadd45b; Cxcr4; Ctnna3; Dapk2; Rac2; Vegfd; Fgf5; Fzd5; Bdkrb2; Il2rg; Frat1; Stat5a; Nkx3-1; Gadd45a; Gng8; Hhip* |
| G alpha (q) signaling events | Reactome | 0.00604 | 0.07505 | *Trh; Bdkrb2; Grp; Tbxa2r; Qrfpr; Npffr1; Trpc6; Grpr; P2ry1; Tacr3; Nts; Gng8; Cysltr1* |
| Basal cell carcinoma | KEGG | 0.00751 | 0.08861 | *Fzd4; Fzd5; Fzd9; Gadd45b; Gadd45a; Hhip* |
| PCP/CE pathway | Reactome | 0.00907 | 0.09929 | *Fzd4; Rac2; Fzd5; Ror1; Ror2* |
| Th17 cell differentiation | KEGG | 0.00926 | 0.09929 | *Rara; Il2rg; Irf4; Nfkbia; Stat5a; Zap70; Lck; Il1r1* |
